# Host microRNA Target Prediction in SARS-CoV-2 and Hepatitis E Virus Genomes: Insights from RNAhybrid Analysis

**DOI:** 10.1101/2025.06.23.661149

**Authors:** Zerin Tasnim, Fariha Islam

## Abstract

MicroRNAs (miRNAs) are key post-transcriptional regulators of gene expression that can influence viral replication and pathogenesis by binding to viral RNA genomes. This study uses RNAhybrid to predict binding sites of three abundant human miRNAs—hsa-miR-155-5p, hsa-miR-21-5p, and hsa-let-7a-5p—within the genomes of SARS-CoV-2 (Bangladesh isolate OM967280.1) and Hepatitis E Virus genotype 1 (NC_001434.1). Our results show stable hybridization events with minimum free energy (mfe) values ranging from −23.8 to −28.1 kcal/mol, indicating strong potential interactions. These findings suggest that host miRNAs may modulate viral RNA stability or translation, impacting viral pathogenesis. Further experimental validation is necessary to confirm these regulatory relationships and explore their therapeutic potential.

## Introduction

MicroRNAs (miRNAs) are short, ~22 nucleotide non-coding RNAs that regulate gene expression by binding to complementary sequences within target RNAs, leading to translational repression or degradation (Bartel, 2004). Beyond endogenous gene regulation, miRNAs have been implicated in host-virus interactions, where they can target viral genomes or transcripts to modulate infection outcomes (Jopling, 2005; Skalsky & Cullen, 2010).

SARS-CoV-2, the causative agent of COVID-19, is an RNA virus whose interaction with host miRNAs is an active area of research, as these interactions may influence viral replication or immune evasion (Fulzele et al., 2020). Similarly, Hepatitis E Virus (HEV), another positive-sense RNA virus, may also be subject to miRNA regulation, although data remain limited (Sayed et al., 2017).

Here, we computationally predicted putative binding sites of three human miRNAs—hsa-miR-155-5p, hsa-miR-21-5p, and hsa-let-7a-5p—on SARS-CoV-2 and HEV genomes using the RNAhybrid tool (Rehmsmeier et al., 2004). These miRNAs were selected for their known roles in antiviral immune responses and inflammation (O’Connell et al., 2012; Sheedy, 2015).

## Materials and Methods

### Sequence Data

Complete genome sequences for SARS-CoV-2 Bangladesh isolate (OM967280.1) and Hepatitis E Virus genotype 1 (NC_001434.1) were retrieved from the NCBI GenBank database.

### miRNA Sequences

Human miRNA sequences for hsa-miR-155-5p, hsa-miR-21-5p, and hsa-let-7a-5p were obtained from miRBase (Kozomara et al., 2019).

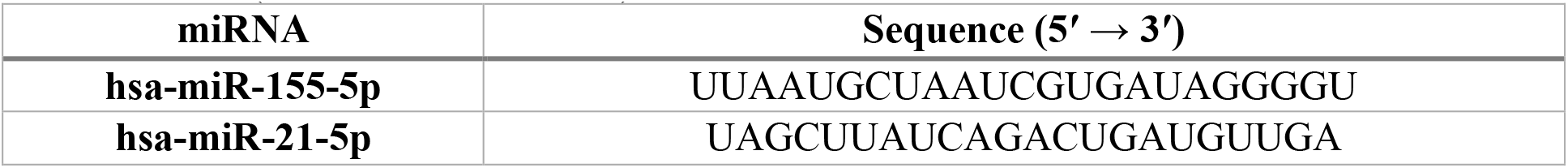
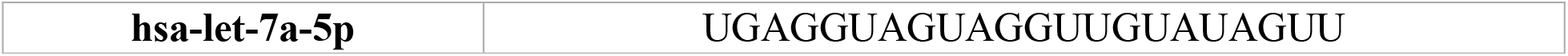

### miRNA Target Prediction

RNAhybrid (version 2.2) was employed to predict miRNA binding sites on the viral genomes (Rehmsmeier et al., 2004). The tool computes minimum free energy (mfe) hybridizations, considering perfect or near-perfect seed pairing and allowing G:U wobble pairs. Parameters were set with an energy threshold of −20 kcal/mol to filter biologically relevant interactions.

## Results

Multiple potential binding sites for the three miRNAs were identified on both viral genomes. The most stable binding energies for each miRNA on SARS-CoV-2 and HEV are summarized in Table 1.

**Table 1.** Predicted microRNA Binding Sites on Viral Genomes.

| miRNA | Binding Position | mfe (kcal/mol) | Target Virus | Accession | Notes |
| --- | --- | --- | --- | --- | --- |
| **hsa-miR-155-5p** | 6619 | $-24.8$ | SARS-CoV-2 BD | OM967280.1 | Strong seed pairing |
| **hsa-miR-21-5p** | 9388 | $-24.8$ | SARS-CoV-2 BD | OM967280.1 | Moderate stability |
| **hsa-let-7a-5p** | 7087 | $-27.6$ | SARS-CoV-2 BD | OM967280.1 | Most stable binding |
| **hsa-miR-155-5p** | 3762 | $-28.1$ | HEV genotype 1 | NC_001434.1 | Strong binding site |
| **hsa-miR-21-5p** | 3729 | $-23.8$ | HEV genotype 1 | NC_001434.1 | Moderate binding |
| **hsa-let-7a-5p** | 5522 | $-25.8$ | HEV genotype 1 | NC_001434.1 | Stable binding |

**Figure 1.**
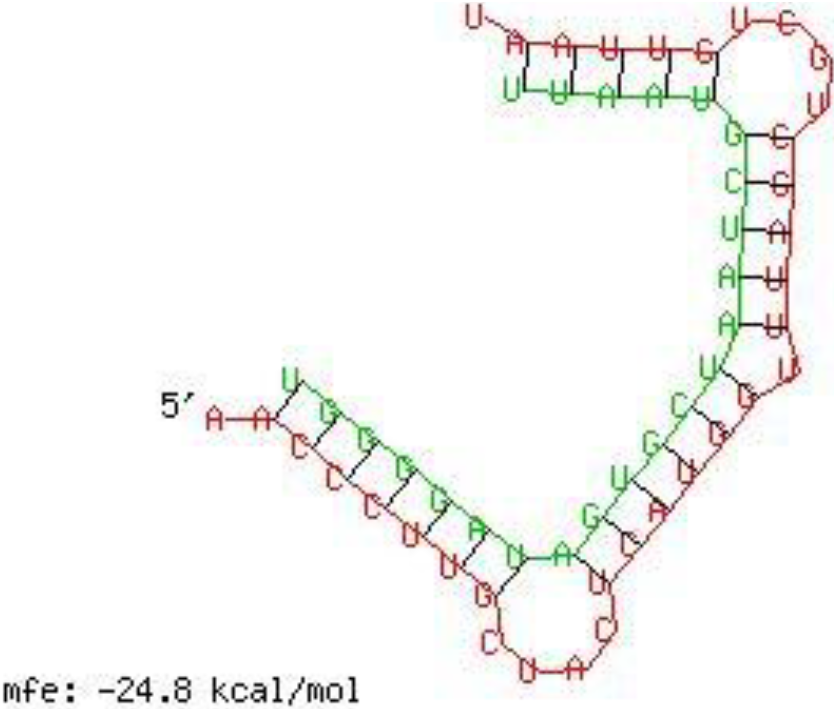
Predicted hybridization between human microRNA hsa-miR-155-5p and the SARS-CoV-2 Bangladesh isolate (OM967280.1)

**Figure 2.**
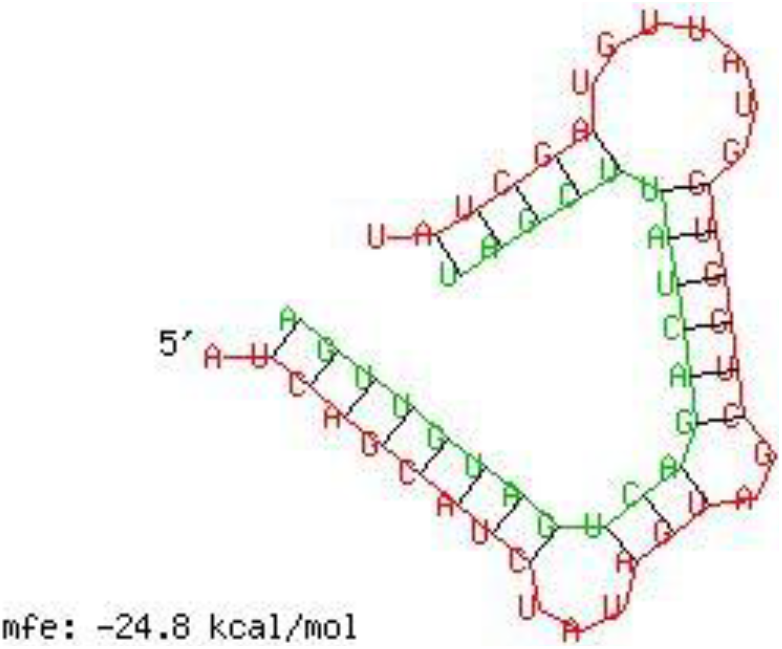
Predicted interaction between hsa-miR-21-5p and SARS-CoV-2 Bangladesh isolate (OM967280.1)

**Figure 3.**
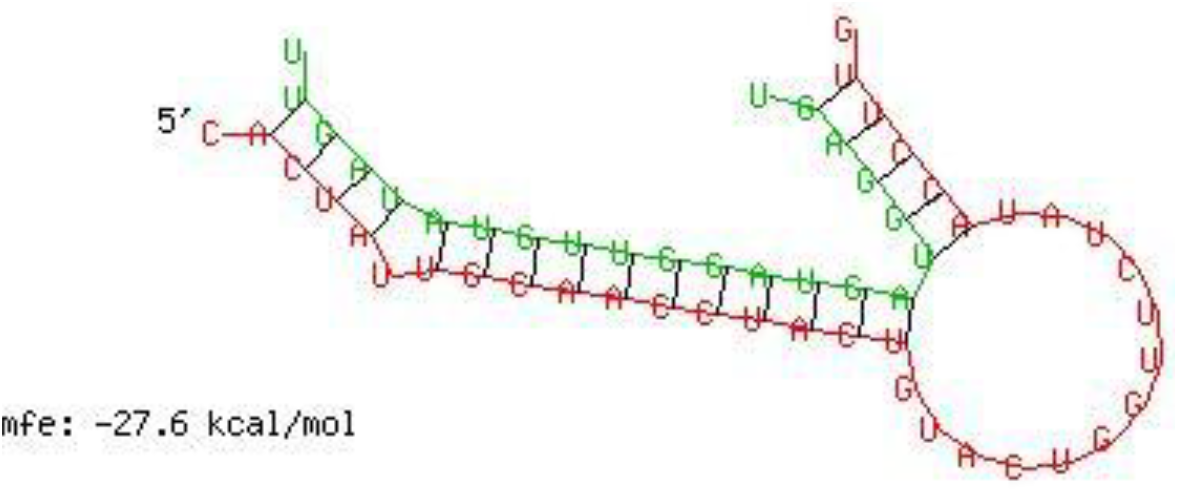
Predicted interaction between hsa-let-7a-5p and SARS-CoV-2 Bangladesh isolate (OM967280.1)

**Figure 4.**
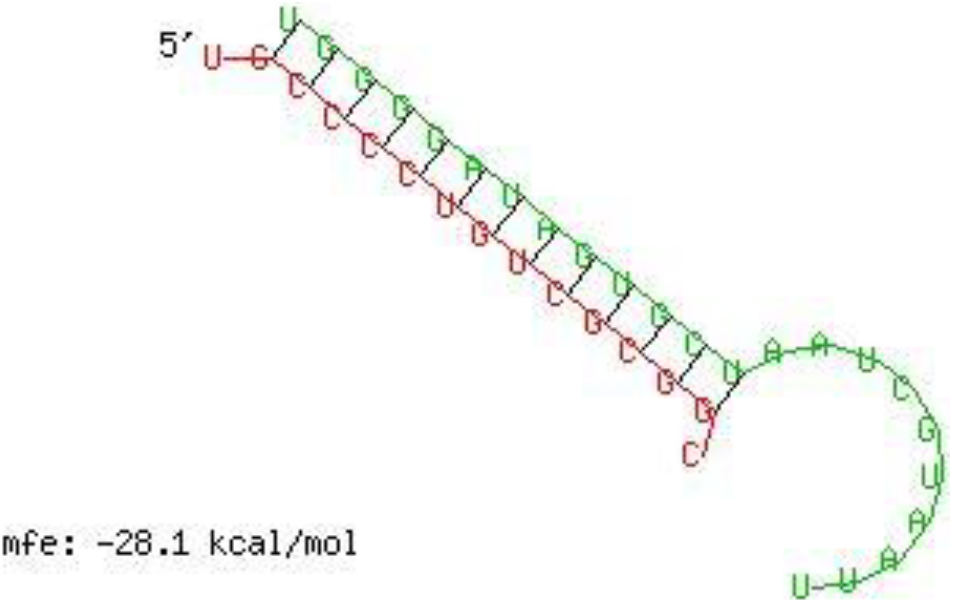
Predicted binding of hsa-miR-155-5p to Hepatitis E Virus genotype 1 (NC_001434.1) genome at position 3762.

**Figure 5.**
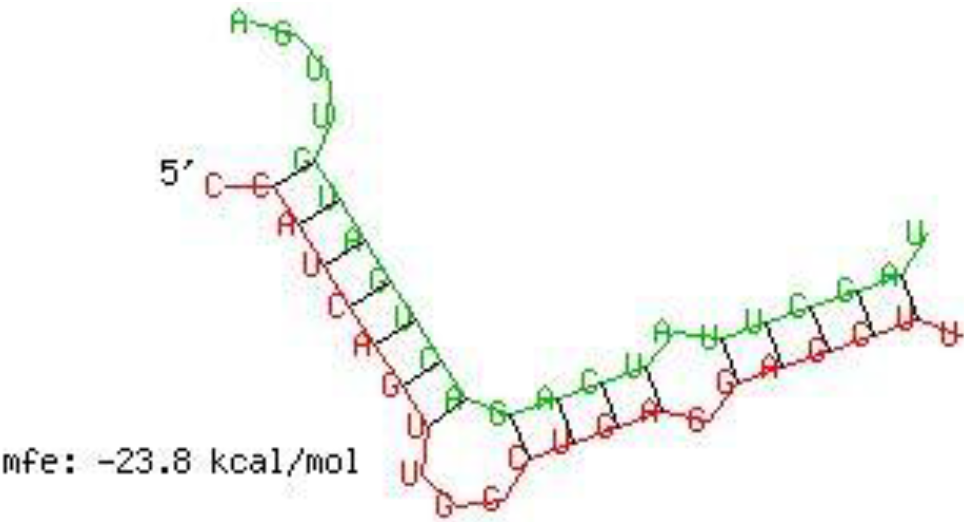
Predicted binding of hsa-miR-21-5p to Hepatitis E Virus genotype 1 (NC_001434.1) genome at position 3729.

**Figure 6.**
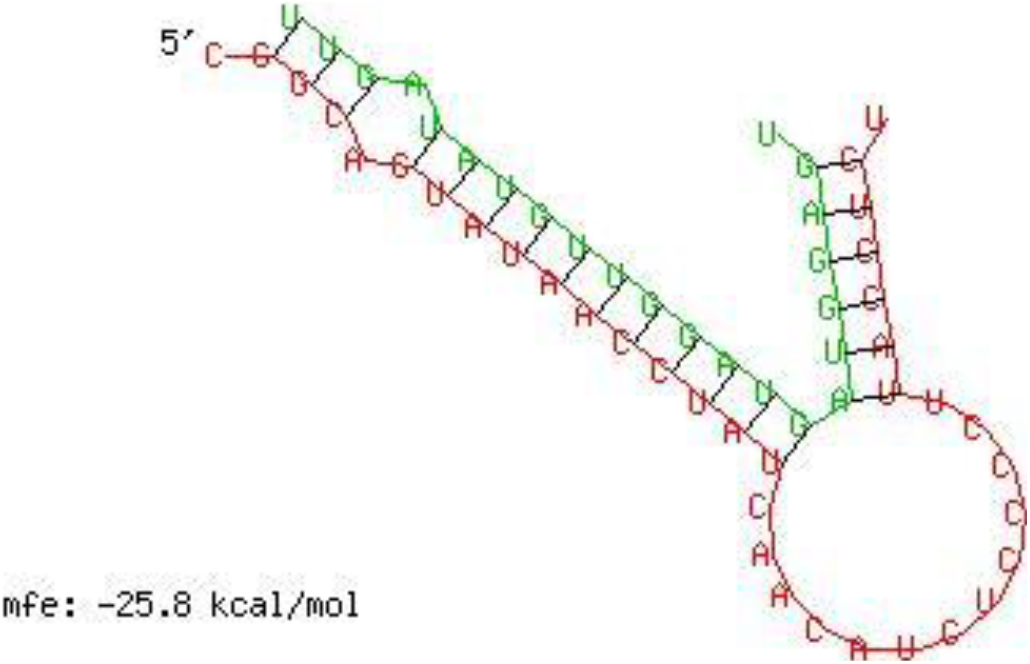
Predicted binding of hsa-let-7a-5p to Hepatitis E Virus genotype 1 (NC_001434.1) genome at position 5522.

The strongest predicted interaction was between hsa-let-7a-5p and SARS-CoV-2 at nucleotide position 7087, with an mfe of −27.6 kcal/mol. The stable seed pairing suggests functional binding, which may impact viral RNA stability or translation. Similarly, hsa-miR-155-5p exhibited strong binding to HEV at position 3762 (mfe −28.1 kcal/mol)

## Discussion

Our computational predictions provide evidence for potential host miRNA targeting of SARS-CoV-2 and HEV genomes. The identified binding sites correspond to stable miRNA–viral RNA duplexes with energetics consistent with functional interactions reported in other RNA viruses (Luna et al., 2015). For example, hsa-miR-155 has been previously linked to antiviral immune modulation (O’Connell et al., 2012), and hsa-let-7a regulates inflammatory pathways (Schulte et al., 2013).

Experimental validation using reporter assays and infection models will be essential to confirm the regulatory roles of these miRNAs. If confirmed, these miRNAs could serve as therapeutic targets or biomarkers for viral infections.

## Conclusion

The predicted miRNA binding sites in SARS-CoV-2 and HEV genomes suggest a potential layer of host post-transcriptional regulation during infection. This study highlights hsa-let-7a-5p and hsa-miR-155-5p as candidates for further investigation into antiviral host defense mechanisms.

